# Cytoskeleton mechanics determine resting size and activation dynamics of platelets

**DOI:** 10.1101/413377

**Authors:** Aastha Mathur, Sandra Raquel Correia, Serge Dmitrieff, Romain Gibeaux, Iana Kalinina, Tooba Quidwai, Jonas Ries, Francois Nedelec

## Abstract

Platelets are cell fragments of various size that help maintain hemostasis. The way platelets respond during a clotting process is known to depend on their size, with important physiological consequences. We characterized the cytoskeleton of platelets as a function of their size. In resting Human and Mice platelets, we find a quadradic law between the size of a platelet and the amount of microtubule polymer it contains. We further estimate the length and number of microtubules in the marginal band using Electron and Super-resolution microscopy. In platelets activated with ADP, the marginal band coils as a consequence of cortical contraction driven by actin. We observe that this elastic coiling response is accompanied by a reversible shortening of the marginal band. Moreover, larger platelets have a higher propensity to coil. These results establish the dynamic equilibrium that is responsible for platelet size and differential response on a more quantitative level.

**Highlights:**

- Platelet size scales consistently with amount of polymerized tubulin in both mouse and human.
- Polymerized actin is required for ADP-induced marginal band coiling.
- Upon activation, the marginal band exhibits a reversible visco-elastic response involving shortening.
- Larger marginal bands have a higher propensity to coil than shorter ones.

**In brief:** The cytoskeleton is adapted to platelet size and its mechanical properties determine propensity of a platelet to undergo morphological changes in response to agonists.

## Introduction

Blood clotting involves a complex interplay of cascading chemical reactions that influence the fluid and cellular components of blood to form a clot. In mammals, platelets, which are 2-3µm sized, discoid cell fragments, are an essential component of the haemostatic machinery. Formed by fragmentation of the rare, large (10-65µm {Levine:1982vu}) and polyploidy bone marrow residents called megakaryocytes; platelets themselves are relatively simple in shape and internal architecture (Figure 1A). To fulfill their function of forming a blood clot, these small cytoplasmic fragments are able to undergo complex morphological changes during platelet activation process. The first step of this activation sees the cell going from a flat, disc-like shape to a more spherical shape from which protrusions appear, and is thus called the disc-to-sphere transition. Activation of platelets can be triggered by multiple factors including soluble agonists like adenosine-di-phosphate (ADP), thrombin, etc. that induce the disc-to-sphere transition while platelets are still in solution. ADP is considered a weak agonist of platelet activation, as physiologically, it is often present only in addition to other agonists to form a clot.

**Figure 1:**
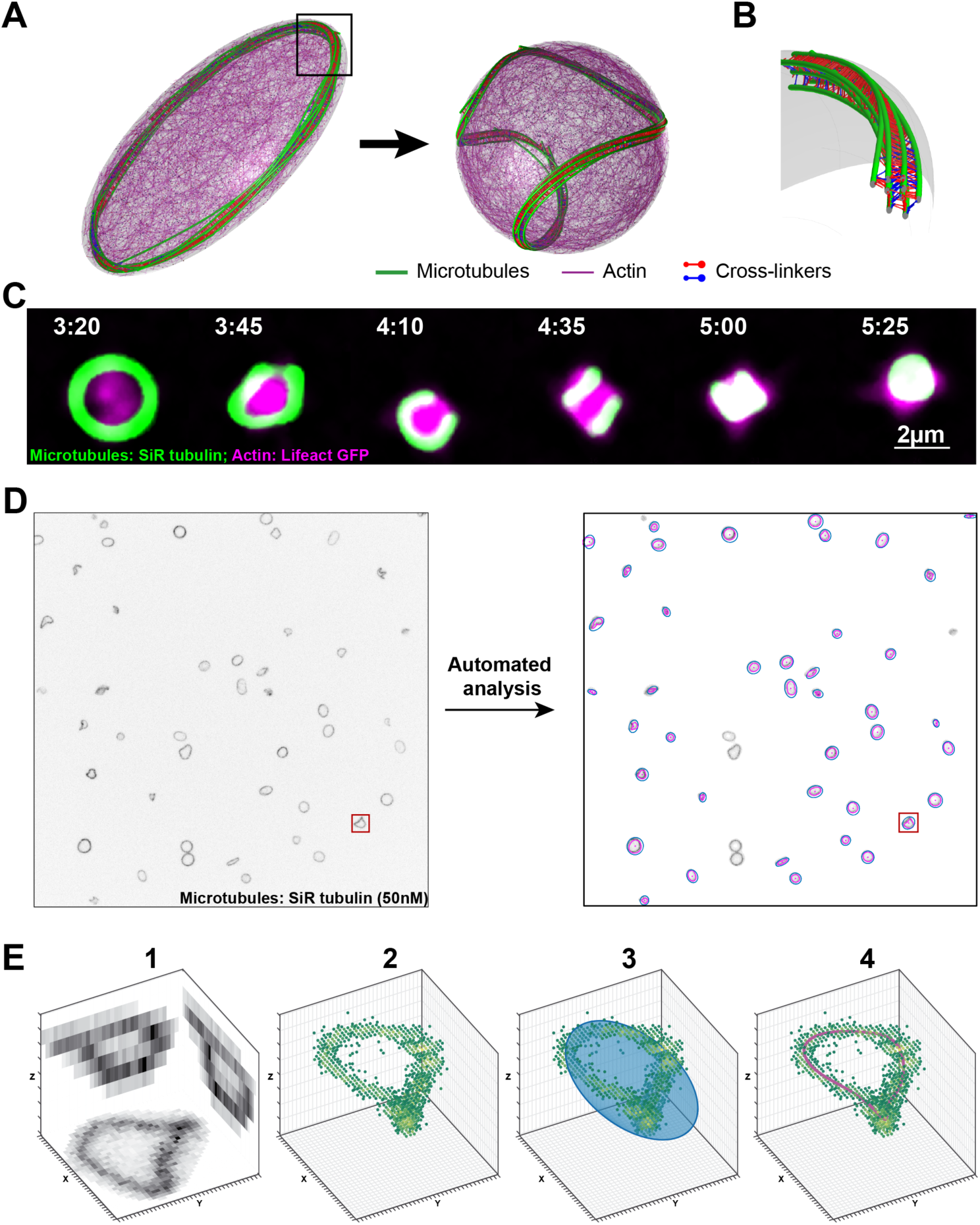
Quantifying marginal band morphology and coiling dynamics. **(A)** Schematic representation of cytoskeletal organization in platelets in discoid and spheroid morphology. **(B)** Zoomed in view of the platelet marginal band (MB) showing microtubules held together by cross-linkers and motors. **(C)** Visualization of cytoskeleton during platelet morphological changes in response to ADP (adenosine-di-phosphate). MB consisting of microtubules undergoes coiling while actin polymerizes and localizes cortically. Time in min:sec since start of flow of ADP containing solution. **(D)** Typical microscopic field of view (100µm x 100µm) showing marginal bands after 200sec of ADP treatment before (left) and after (right) quantification. Blue: fitted ellipse to estimate platelet size; Magenta: fitted 3D curve through MB pixels; Red box: specimen platelet described in E. **(E)** Quantitative parameters describing MB morphology. **1:** XY, XZ and YZ projections of single MB. **2:** Isolated MB voxels color-coded for intensity. Summed intensity estimates amount of polymerized tubulin. **3:** 2D plane (Blue) is fitted to MB voxels so as to minimize deviations orthogonal to the plane. Blue ellipse is fitted to estimate platelet size. **4:** A closed 3D curve (magenta) is fitted to pixels weighted by intensity. This gives length of the MB. The degree of coiling is determined based on average distance of the magenta curve from the fitted plane. All microscopy images are maximum intensity projections.

The cytoskeleton plays an important role in various aspects of platelet activity, from their biogenesis, to activation, to clot formation and eventual clot retraction. Mechanical properties of the cytoskeleton are implicated in formation, extension and fragmentation of platelet precursors called pro-platelets, finally affecting the equilibrium size of a resulting platelet (Thon et al., 2012). The mean platelet volume (MPV) is an important diagnostic measure, as many bleeding disorders are known to be associated with abnormal platelet size and morphology. Many giant platelet disorders are associated with mutations in non-muscle myosin heavy chain 9 ({Seri:2000dl} {Kelley:2000ff} {Zhang:2012vg}) that affect platelet contractility. Other giant platelets from Bernard-Soulier syndrome models show altered microtubule cytoskeleton {Strassel:2009ec}. The contractile forces exerted by myosin on actin {Leon:2007df} {Zhang:2012vg} {Spinler:2015dk}, and pushing forces of sliding microtubules by dynein {Bender:2015jk} are both required for proper pro-platelet elaboration and formation of platelets in correct size and number. Thus, cytoskeletal organization and mechanics is deemed important for maintaining a physiological platelet activity.

In a resting or discoid platelet, microtubules form a closed circular bundle running near the cell periphery, known as marginal band (MB). Early electron microscopy indicated that this ring was made of a single of microtubule coiled over multiple turns (Kowit, Linck, & Kenney, 1988). However, using a microtubule plus-end binding marker, it was later shown to contain 7 to 12 growing microtubules {PatelHett:2008jz}, orientated in both directions equally. Microtubules of the MB are cross-linked {ZuckerFranklin:1969ez} forming a rigid bundle that gives resting platelets their characteristic discoid shape. A loss of microtubules by chemical agents like vincristine or by cold treatment causes platelets to adopt a spherical morphology (White & Rao, 1998).

In addition to the MB, platelets also have membrane associated cortical cytoskeleton containing cross-linked spectrin and actin fibers {Nakata:1987dc} (Hartwig & DeSisto, 1991). The interconnected fibers along with myosin form a 50 to 100µm thick network of mixed polarity lining the inner surface of the plasma membrane. A non-cortical actomyosin meshwork is also present in the bulk cytoplasm, associated to platelet internal membrane structures (Boyles, Fox, Phillips, & Stenberg, 1985). In a resting platelet, polymerized actin accounts for only 40% of platelet actin, as the majority of the G-actin (300-350µM; {Fox:1984wi}) is sequestered or maintained in a soluble form by barbed end capping and filament stabilization {Bearer:2002uu}. During the disc-to sphere transition, this reserve actin assembles and localizes in the cortical region, while MB coils in the characteristic ‘baseball seam curve’. Although this process is correlated with depolymerization of microtubules {Paul:2003wb}, the depolymerization itself is not essential for the disc-to-sphere transition {Shiba:1988tb}. It has been previously reported that molecular motors like myosin, associated with actin {Paul:2003wb} (Hartwig & DeSisto, 1991), and cytoplasmic dynein, associated with microtubules {Rothwell:1997uy}, change their localization and phosphorylation state with ADP induced activation. The chain of events leading to MB coiling remains controversial, with two possibilities being debated. The first one is that activation induces cortical contraction leading to mechanical compression of the MB. In this scenario, the MB initially behaves like a passive elastic coil constrained by the actin cortex (Dmitrieff, Alsina, Mathur, & Nedelec, 2017). Alternatively, platelet activation would activate some microtubule associated motors, possibly kinesins acting within the ring, or dynein molecules anchored at the cortex. The activity of these motors would induce MB lengthening, leading it to coil as the cell confinement remains (Diagouraga et al., 2014). The key observation supporting this second scenario is based on population average of fixed cells: coiled MBs were found to be longer than flat, resting, ones (Diagouraga et al., 2014). To clarify these important mechanistic aspects of platelet response, it seemed essential to assess the evolution of the MB in live cells.

In this study, we systematically quantified the quantity of polymerized microtubules in a population containing thousands of resting platelets, to uncover how this quantity varies with platelet size. By monitoring live platelets in suspension within a microfluidic setup, we visualize and quantify the morphology and length of the MB during the disc-to-sphere transition following activation. This single-cell level study uncovers a size-based heterogeneity in platelet response. Using electron tomography and super-resolution microscopy we further characterize the structure of MB in resting platelets, leading to a detailed characterization of the microtubule cytoskeleton as a function of platelet size.

## Results

### An integrated platform to analyze platelet activation

The disc-to-sphere transition occurs while platelets are in the blood stream, that is, suspended in fluid medium. To image the MB close to these conditions, we developed the diffusion based microfluidic device described in figure S1. This device uses pillar arrays to restrict flows, enabling us to acquire 3D images of platelets in suspension while tracking individual cells. as well as expose them to diffusible molecules, particularly cytoskeleton drugs and platelets agonists such as ADP. We first characterized the time needed for an inert fluorescent tracer, fluorescein, to reach the imaging area (Figure 1B; Supp. Figure 1B) by travelling through the pillared arrays (Supp. figure 1A, P2 and P2). Upon activation by ADP, the platelet cytoskeleton can be observed to reorganize, concomitant with the disc-to-sphere transition in platelet morphology. The shape change occurs within 1 min of ADP treatment. The microtubule MB undergoes coiling while actin polymerizes and localizes in the platelet cortical region, where it can be observed to extend filopodia (figure 1C).

The imaging setup used permits large fields of views (100µm×100µm) containing multiple platelets to be recorded and treated with soluble molecules during imaging (figure 1D). To have access to the population-level and single-cell-level response of platelets, we developed a fully automated image analysis pipeline, able to track individual cells and analyze their time-dependent behavior. To image microtubules, we used 50nM of SiR-tubulin, a low dose that is thought to have little physiological effects, and summed the intensities of the pixels belonging to the MB. This is a proxy for the amount of polymerized tubulin as SiR-tubulin ((Lukinavičius et al., 2014)) only binds polymerized tubulin (figure 1 E2). In brief, the image is segmented, platelets identified and a closed curve is fitted to the intensity-weighted pixels of each MB in 3D (figure 1 E4), the contour of which gives the MB length. We also determine a plane that minimizes all orthogonal distances to the voxels identified as originating from the MB (figure 1 E3). A flat MB in a discoid platelet lies within this plane while a coiled MB in a spherical platelet bends out of it. The average normalized distance from the fitted curve to that plane gives a “degree of coiling” of the MB. Together with the contour length, this dimension-free quantity allows us to characterize the key aspects of MB evolution. This setup allowed us to query a large population of resting platelets and to quantitatively measure changes in MB morphology of individually tracked platelets during the course of activation.

### Marginal band size varies in a platelet population and scales with the polymerized tubulin content

Platelets naturally vary in size possibly due to their formation process, age, contents, etc. Dispersion of size can strongly affect both the chemical and the mechanical response of platelets. Such variations indeed affect platelet activity, making their size an important diagnostic measure {Frojmovic:1982wk}. With our light microscopy setup, we observed a considerable natural variation in resting MB sizes as well as intensities (figure 2A). Large samples of resting mouse platelets (n=6358 platelets from 13 experiments) reveal however a consistent distribution, with a MB length of 8.98±1.44 µm (mean and SD, figure 2B). Next, to uncover how the cytoskeleton determines the size of platelets, we measured the relationship between MB length (*L*_*MB*_) or radius (*R* = *L*_*MB*_/2π) and the amount of polymerized tubulin (*I*_*MT*_), estimated by the total background-subtracted fluorescence intensity of the MB. The best fit is obtained for *I*_*MT*_ ∝ *R*^*α*^ with *α* = 1.93 ±0.035, giving an approximate power dependence *I*_*MT*_ ∝ *R*^2^ (figure 2C) (see supplementary material for the regression procedure).

**Figure 2:**
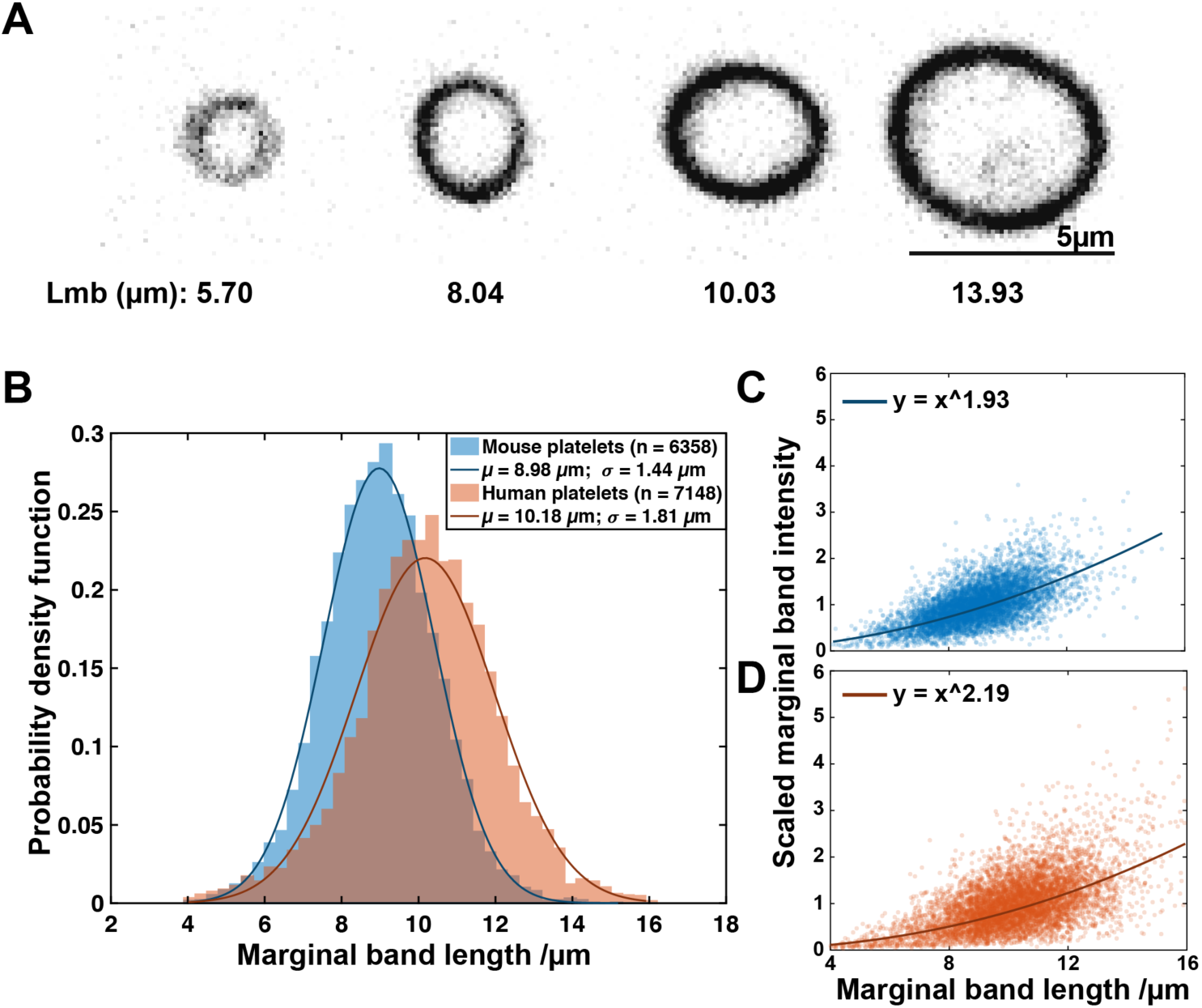
Population variation in MB length and the underlying scaling with polymerized tubulin content. **(A)** Maximum intensity projections of isolated mouse platelet MBs of different sizes from the same field of view. Intensity corresponds to SiR tubulin fluorescence. Marginal band length is noted below the images. **(B)** Histogram of MB length distribution in mouse platelets (mean: 8.98µm; standard deviation: 1.44 µm; number of platelets: 6358; number of experiments: 13) and human platelets (mean: 10.17µm; standard deviation: 1.81 µm; number of platelets: 7148; number of experiments: 5). **(C)** Scaling relationship between MB length and SiR tubulin intensity for mouse platelet population in fig. 2B. SiR tubulin intensity, used as a proxy for polymerized tubulin mass, increases as a power of 1.93±0.035 of MB length. **(D)** MB length and SiR tubulin intensity scaling for human platelet population in fig. 2B. Human platelet sizes vary as a power of 2.19±0.039 with the amount of polymerized tubulin.

Curious to know if this law would hold true for other species, we turned to human platelets, which are approximately twice more voluminous than their mouse counterparts ({Schmitt:2001eb}). Using the same methods, we measured a large sample of human platelets (n = 7148). Their MB length was 10.18±1.81µm (mean and SD, figure 2B). On an average, human platelets were larger than mouse platelets (two-sampled t-test; p<2.2e-16) as expected. Upon measuring the dependence of MB length on microtubule intensity we found the power exponent for human platelets to be around 2.19 ±0.039 (figure 2D). Thus, both mouse and human platelets follow approximately *I*_*MT*_ ∝ *R*^2^. A similar power law underlying the scaling in both mouse and human platelets alludes to the presence of common mechanisms that control platelet sizes in these organisms.

### The platelet cytoskeleton reorganizes visco-elastically in response to activation with ADP

With the aim of understanding the mechanics of platelet shape change in response to ADP, we first quantified the dynamics in a control sample, measuring the degree of coiling and MB length over time after treatment with ADP. We used the time at which the intensity of the fluorescein tracer reaches 10% of its maximum to align the tracks from different experiments (average shown in figure 3A). This time marked as zero. We found that after a typical lag time, MBs undergo a steep rise in degree of coiling (figure 3B) reaching a peak at time zero before relaxing, suggesting that most of the MBs return to a flat state after a transient coiling. Concomitantly, we measure a reduction in MB length that is maximal around time zero, and followed by a gradual recovery, which is partial and plateaus after 700sec (figure 3C). To ascertain that potential flows and DMSO alone do not cause a change in marginal band morphology, we performed the same experiment without ADP and observed no response (figure 3 B and C). This partially reversible change in platelet morphology is in agreement with the known a response to a mild ADP treatment {Milton:1980vx}.

**Figure 3:**
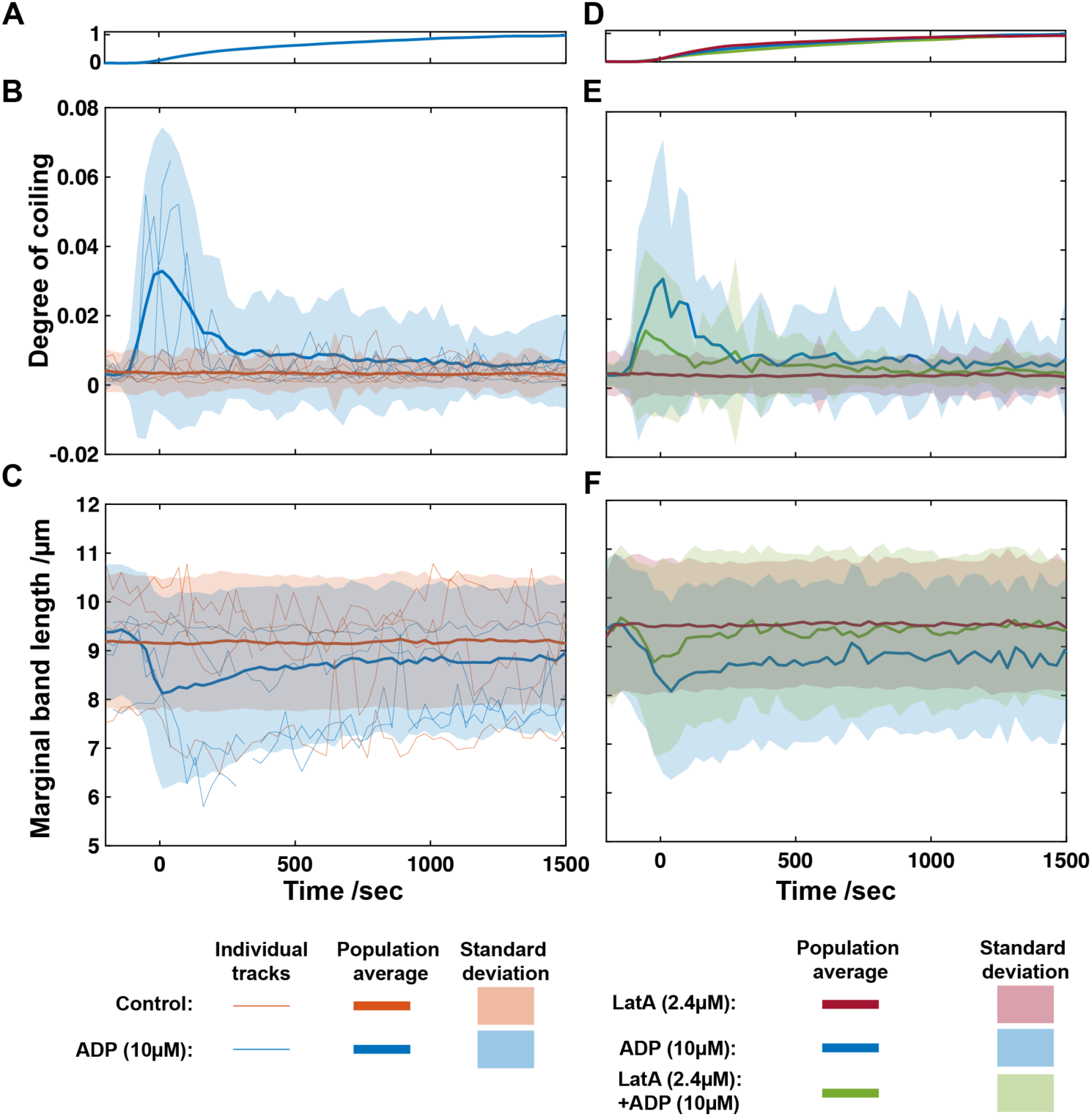
Coiling dynamics of MB population in response to activation with ADP. **(A)** Average intensity of fluorescent tracker molecule, fluorescein (1µM), over time for experiments with ADP treatment. The scaled trace is an average of 13 individual experiments aligned with respect to 10% increase in intensity (Supp. figure 1C and 1D) which is marked as time zero (t = 0). **(B)** Measure of degree of coiling over time for platelet population treated with ADP (Blue; number of tracks: 175; number of experiments: 13) and control platelets subjected to flow with DMSO (Orange; number of tracks: 358; number of experiments: 13). ADP treated MBs undergo a sharp increase followed by relaxation in degree of coiling while controls show no change. **(C)** Change in marginal band length over time for the same platelet population as in 3B. ADP treated MBs undergo a decrease followed by partial relaxation of marginal band length while controls show no change. **(D)** Average intensity of fluorescent tracker molecule, fluorescein (1µM), over time for experiments with ADP only (Blue; n = 4; subset of data in 3A), Lat A and ADP co-treatment (Green; n = 4) and LatA pre-treatment followed by ADP (Red; n = 8). The scaled trace is an average of individual experiments aligned with respect to 10% increase in intensity which is marked as time zero (t = 0). Visibility of the three separate curves might be obscured due to very reproducible dynamics. **(E)** Measure of degree of coiling over time for platelet population treated with ADP only (Blue; number of tracks: 50; number of experiments: 4; subset of data in 3B), Lat A and ADP co-treatment (Green; number of tracks: 33; number of experiments: 4) and LatA pre-treatment and ADP (Red; number of tracks: 230; number of experiments: 8). LatA pretreated platelets show no change in degree of coiling. **(F)** Measure of marginal band length over time for the same platelet population as in 3E. LatA pretreated platelets show no change in MB length.

### Polymerized actin is required to trigger marginal band coiling in response to ADP

In many cellular systems actin organizes into networks able to generate contractile forces. The tension produced by a cortex is typically in the order of 100 pN/µm, which is sufficient to induce coiling. In platelets, the actin cytoskeleton is deemed essential for adhesion, spreading, as well as in motility ({Gaertner:2017hf} (“Direct characterization of cytoskeletal reorganization during blood platelet spreading.,” 2018)). Disrupting polymerized actin was shown to prevent platelet morphological changes {Severin:2012dx}). However, its role in triggering platelet disc-to-sphere transition is disputed (Diagouraga et al., 2014)). To investigate this, we pre-treated platelets with latrunculin A (LatA), which sequesters actin monomers causing depolymerization of actin filaments, and followed it by an ADP treatment (figure 3D). In agreement with previous reports, we confirmed the absence of any coiling of MB when platelets are pre-treated with LatA and then exposed to ADP (figure 3E). Additionally, we measure no reduction in length (figure 3F), suggesting that the shortening of the MB is a consequence of the force applied by actin, although we can only conclude that presence of polymerized actin is necessary to induce marginal band coiling and shortening in response to ADP based activation.

### Actin perturbation reveals mechanical properties of marginal band

We have shown that a population of platelet marginal bands responds to actin-driven morphology changes by an initial quick elastic coiling and a slower viscous rearrangement. We also showed that the initial coiling is triggered by actin contraction. To address the role of actin in viscous rearrangement of the MB, we depolymerized actin during MB coiling. This was executed by simultaneous treatment of platelets with a solution containing both ADP and LatA. In response to LatA and ADP co-treatment, we see an initial increase in degree of coiling with dynamics similar to the control cases, but up to a lower extent. This is followed by a slower relaxation to flat marginal band morphology (figure 3E). At the same time, we find that the marginal band length decreases initially, like controls but to a lower extent, and then recovers to near initial values, in contrast with ADP-treated platelets in the absence of LatA (figure 3F). Thus, in absence of sustained actin contraction, marginal band length changes are reversible. This data confirms that polymerized actin is required for length reduction of MB, as its dynamic depolymerization leads to a reversible response. Additionally, it shows that viscous rearrangement of the MB leading to its length reduction is induced in response to sustained actin contraction.

### Actin compresses the Marginal Band in resting platelets

After establishing that actin contraction in activated platelets causes MB coiling and reduction in length, we wondered if the length of the MB in resting platelets depended on cortical tension, as one would expect from the balance of force between the marginal band extension and the tension of the cortex, containing actin and spectrin networks. For this purpose, we tracked individual platelets live for 30 minutes after LatA treatment. On a population level MB length does not change appreciably. However, when we binned platelets by their initial lengths (figure 4A inset), we found that the shorter MBs showed, a significant (Welch Two Sample t-test p-value: 0.0082), up to 10% increase in length, while they remained uncoiled (figure 4A). Longer MBs remain mostly unchanged (figure 4B). This data shows that actin is compressing the MB, having an effect that is measurable in small platelets. However, even when there is an effect, LatA only induces a moderate elongation of the MB, suggesting that another cellular structure that is not made of actin is restraining the MB, perhaps the spectrin cortex.

**Figure 4:**
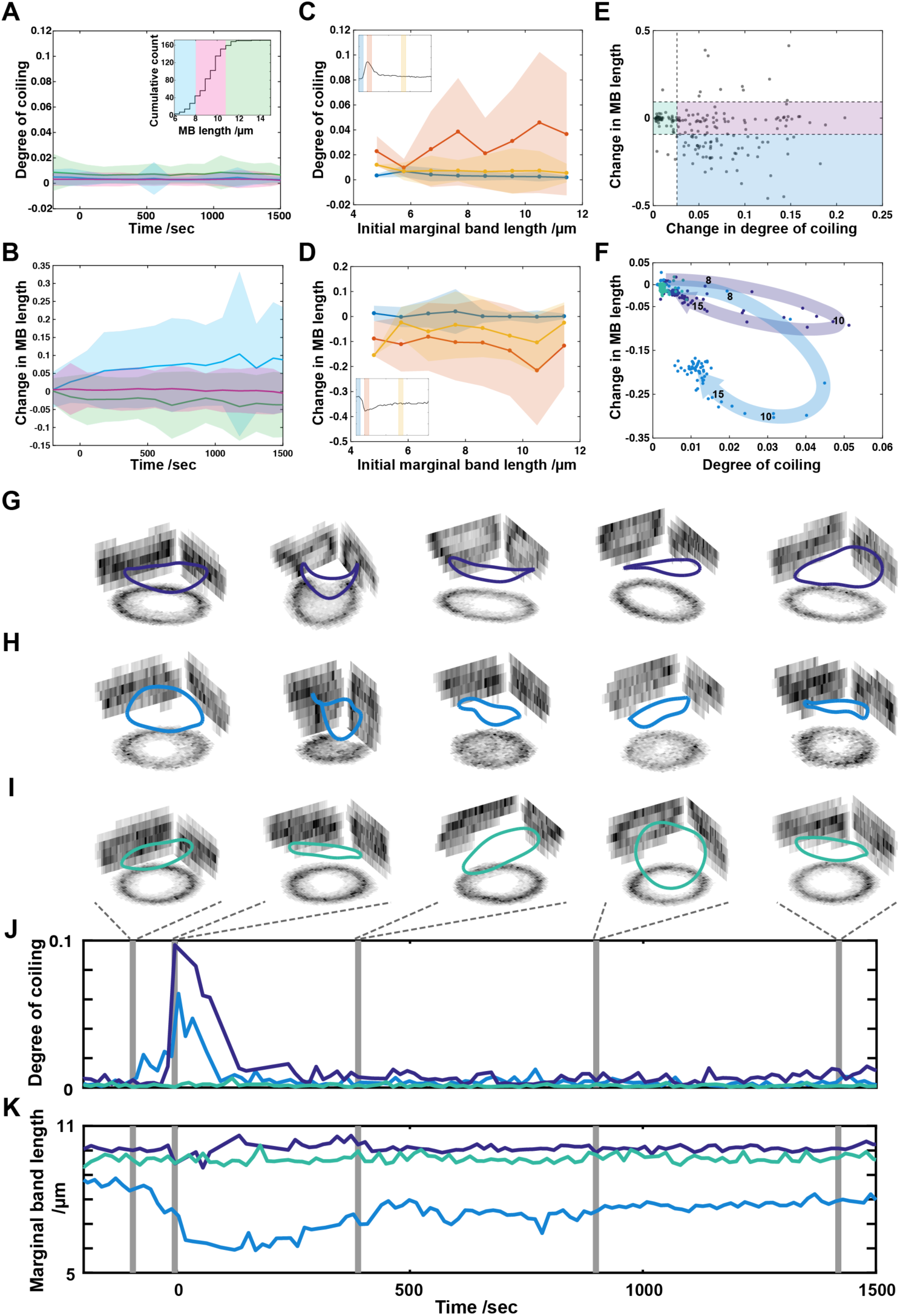
Mechanics of MB coiling response. **(A)** Change in degree of coiling of MBs over time upon LatA treatment. The different colours represent platelets binned by initial MB length, as marked in the inset, which shows the cumulative count of platelets (n: 172; number of experiments: 8). **(B)** Change in MB length over time upon LatA treatment. Colours correspond to figure 4A inset. **(C)** Change in degree of coiling of MBs upon ADP treatment as a function of initial length. The dataset is same as in 3B. The different colours are different time bins as marked in the inset, which shows the degree of coiling over time for ADP treatment. **(D)** Change in length of MBs upon ADP treatment as a function of initial length. The dataset is same as in 3C. The different colours are different time bins as marked in the inset, which shows the length of MB over time for ADP treatment. **(E)** Identification of platelet response subcategories based on changes in marginal band length and degree of coiling. Maximum change in degree of coiling gives elastic buckling response, plotted on x-axis, and relative change in between initial and final length of MB gives viscous rearrangement response, plotted on y-axis (n=175). Dashed lines show thresholds used to assign response subcategories as defined by mean ±standard deviation of the control tracks (n=358). The purple area covers elastic response (number of tracks: 59). The blue area covers visco-elastic response (number of tracks: 58). The green area covers unresponsive platelets (number of tracks: 44). **(F)** Average time traces of platelets in different response categories with the elastic response parameter, i.e. degree of coiling, on the x-axis and the viscous response parameter, i.e. change in MB length, on the y-axis. Examples of tracked platelets where MB undergo, **(G)** elastic and **(H)** visco-elastic responses. Marginal band in **(I)** is unresponsive to ADP treatment. Plots show maximum intensity XY, XZ and YZ projections of isolated platelet MBs at different time points as indicated in the plots below. Intensity corresponds to SiR tubulin fluorescence. Colored 3D curves are fitted to pixels to obtain quantitative parameters. **(J)** Degree of coiling over time for tracked platelet marginal bands above. The tracks have an elastic response parameter of 0.09 for elastic, 0.04 for visco-elastic and 0.00 for unresponsive MBs. **(K)** Marginal band length over time for tracked platelets corresponding to 4E, 4F and 4G. The tracks have a viscous response parameter of 0.00 for elastic, −0.11 for visco-elastic and 0.02 for unresponsive MBs.

### Initial length determines marginal band coiling propensity

It has been previously proposed that the size of a platelet is defined by a mechanical equilibrium between the elastic bending forces of the microtubule bundle and cortical tension of membrane associated cytoskeleton ((Thon et al., 2012) (Dmitrieff et al., 2017)).

Our data offers a large heterogeneity in platelet response to ADP and in platelet size. We could therefore analyze the effect of initial size heterogeneity in platelets on their subsequent response to ADP. Having access to individually tracked platelets, we compared their initial and the final state as a function of initial MB length. We found that cells with longer MBs achieve a higher degree of coiling at the peak of response (figure 4A). At longer times, all MBs become flat again. Indeed from purely mechanical arguments (Dmitrieff et al., 2017), one can expect larger MBs to be more prone to coiling than the smaller ones.

Additionally, we found the decrease in MB length is also dependent on their initial length, as MBs that are initially longer also undergo a larger decrease in length (figure 4B). An incomplete recovery of MB length is also evident at longer times. However, the standard deviation of population response is large suggesting that the observable could be a combination of different subcategories.

To identify potential, mechanistically distinct, categories of coiling, we calculated two response parameters for each platelet track. The first response parameter quantified the change in degree of coiling which refers to the elasticity of MB manifested by out of plane buckling. The second response parameter measured a relative decrease in MB length that is indicative of MB viscosity as revealed by its rearrangement. All the recorded tracks were then binned based on thresholds (figure 4C) calculated using control tracks (response parameter population mean ±standard deviation). Finally we obtained two different categories of mechanical responses. The first category comprised platelets that undergo coiling but no significant change in MB length. Such changes are the hallmark of an elastic response. The second category contained platelets that showed both, increase in coiling as well as a decrease in length of MB. These MBs were considered to be responding visco-elastically. A third category comprised of resilient platelets, which show neither coiling nor a length response. These mechanical subcategories follow typical paths as they traverse the parameter space defined by range of values for degree of coiling and MB length over time (figure 4D). The elastically responding platelets move mostly horizontally, while the visco-elastically responding platelets move diagonally, i.e. they change in both, the MB length and degree of coiling. We can observe the changes in MB morphology of some examples of tracked platelets undergoing elastic (figure 4E) and visco-elastic (figure 4F) coiling as well as unresponsive ones (figure 4G). The corresponding degree of coiling and MB length of these platelets can be observed in figure 4H and figure 4I respectively. These different mechanical responses could arise from a difference in initial mechanical properties of the platelet cytoskeleton.

### Structural techniques reveal composition of marginal band

To understand the mechanical response of platelets, we needed to understand the architecture of its marginal band. We therefore turned to electron tomography (EMT) to obtain precise structural information about microtubule organization in the MB. We obtained a dataset consisting of four human platelets marginal bands (figure 5 A to D) in resting condition. This allowed us to count the number of microtubules, track individual microtubules in space, and annotate microtubule ends as having plus or minus polarity based on their morphology ({Hoog:2011cs}, (Gibeaux et al., 2012)). These results indicate that the MB is a bundle of microtubules with mixed polarities, in agreement with the earlier light microscopy observations {PatelHett:2008jz}, but not with the older EM observations (Kowit et al., 1988). We also observed that neither the plus, nor the minus ends of microtubules are clustered, indicating lack of an organizing center. Additionally, we found existence of parallel and anti-parallel microtubule overlaps. These sites, as shown in other systems, can serve as a location for force generation by microtubule-associated motors like kinesin or dynein. We were also able to visualize the coiled state of the marginal band (Supp. figure 2).

**Figure 5:**
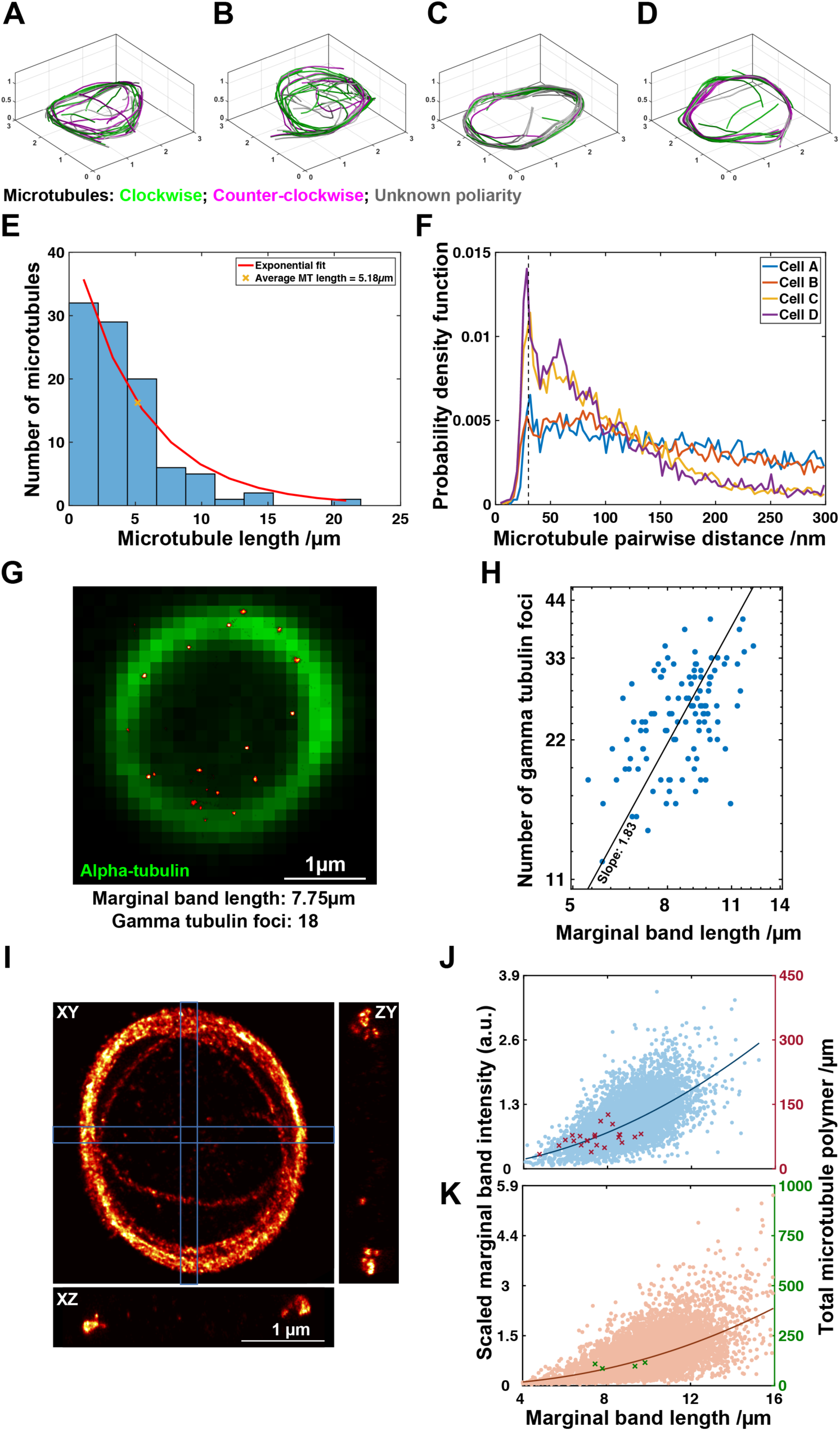
Structural characterization of platelet MB. **(A to D)** Electron tomography dataset showing detailed information about microtubule position and ends within the MB of human platelets. Individual microtubules are color-coded by polarity. This confirms the presence of mixed polarity bundles. **(E)** Microtubule length distribution from all four MBs pooled together is approximated by an exponential function. This yields a characteristic microtubule length of 5.18µm. **(F)** Centre-to-centre pairwise distances between microtubules in four different MBs. The pronounced peak at a distance of 30nm (dashed line) in platelets C and D is reduced in platelets A and B. **(G)** Example image of a diffraction-limited alpha-tubulin labeled MB in green overlaid with super-resolved gamma-tubulin foci (red-hot) marking the microtubule minus ends. **(H)** Scaling between MB length (from diffraction limited alpha tubulin imaging) and number of microtubules (as estimated by number of foci with super-resolution gamma-tubulin imaging), reveals an approximately squared dependence. (I) Example super-resolution microscopy image of MB labeled for alpha tubulin. Central panel shows maximum intensity projection along axial dimension. Side panels show maximum intensity projections along lateral dimensions marked with the blue boxes. Such images are used to estimate the amount of polymerized tubulin in MB as described in Supp. figure 4. **(J)** MB intensity scaling with MB length for mouse platelets calibrated with super-resolution microscopy measurements. Blue data cloud corresponds to the primary y-axis. Data as in plot 2B. Calibration values in red are plotted on the secondary y-axis. **(K)** MB intensity scaling with MB length for human platelets calibrated with electron tomography measurements. Orange data cloud corresponds to the primary y-axis. Data as in plot 2B. Calibration values in green are plotted on the secondary y-axis.

In our tomography dataset, microtubules could be individually followed, which enabled us to extract many quantitative parameters, some of which were previously inaccessible. Firstly, we measured the total length of polymerized MTs, which added up to 101.84μm ±12.63; a quantity in line with pervious estimates ((Kenney & Linck, 1985), {Steiner:1979co}). Secondly, we counted the number of microtubules in each platelet, yielding and average value of 24 ±8.1. This value, although variable, is much larger than previous estimates from marginal band cross-section (8 to 12; {Behnke:1966vc}), or localization of microtubule plus end (EB1: 8.7±2) or minus end (Gamma tubulin: 9.06±1.6) markers ({PatelHett:2008jz}). To identify the distribution of microtubule lengths in platelet population, we pooled data from the four marginal bands. A histogram in figure 5E shows this distribution with an exponential fit that has a characteristic length of 5.18μm. Exponentially distributed lengths are previously shown to be a hallmark of the microtubule population undergoing dynamic instability ((Mitchison & Kirschner, 1984), {Dogterom:1993bo}). The high spatial resolution of this dataset also enabled us to measure pairwise distances between microtubules (figure 5F). This data shows that the MBs that have a more compact morphology (cell3 and cell4), maintain preferred distances between microtubules (peak at 30nm; secondary peak at 60nm), suggesting that these distances are imposed by specific cross-linkers. The more loosely packed MBs (cell1 and cell2) lose the over-representation of the 30nm distance, indicating a loss of potentially specific cross-linkers.

Since MB sizes show high variability, we complemented our EMT data with structural data obtained with localization based super-resolution microscopy (SRM) to estimate the number of microtubules in platelets. Since microtubule minus-ends are known to be less dynamic than the plus-ends in most systems ((Dammermann, Desai, & Oegema, 2003)), we labeled the platelets for gamma-tubulin protein, which localizes to the minus-ends ({Li:1995vz}). Three-dimensional data was recorded and resolved to identify distinct localization sites (figure 5G) that were used as a proxy to count the minus ends and, hence, the number of microtubules. We found that, on an average, platelets contain 25.2 ±6 foci per platelet, which matches the number of microtubules we measured with EMT. We also looked at how the number of microtubule in platelets scales with MB length. For this, we recorded images of the same MBs in SRM for gamma tubulin and diffraction limited immune-fluorescence imaging for alpha tubulin. We were able to measure both, the number of gamma-tubulin foci and the length of the marginal band, in the same platelet. We found that the power law governing the dependence of number of microtubules in platelet MBs, *N*, to their size, *R*, follows a power law of factor 1.83 (Figure 5H), resulting in the relationship of *N* ∝ *R*^2^. This scaling was predicted based on the mechanics of platelet formation (Thon et al., 2012).

In addition to number of microtubules, we used SRM to estimate the amount of polymerized tubulin in platelets. To this end, we fixed mouse platelets and labeled MBs with anti-tubulin primary and dye-coupled secondary antibodies. Due to the tight bundling of microtubules in the marginal band, we were unable to resolve individual microtubules (figure 5I). However, we used the single microtubules often found in the interior or the exterior of the platelets, to calibrate the fluorescence signal (Supp. figure 4). Based on such calibration, we estimated the total amount of polymerized tubulin as well as MB length for each platelet.

Our larger dataset was imaged with fluorescent light microscopy and it was not possible to calibrate the fluorescence intensity absolutely. However, since we measured the MB size of similar cells with SRM and EMT (figure 2C and D), we were able to calibrate the fluorescence dataset a posteriori. In practice, we calculated a scale for the fluorescence intensity by minimizing the squared distance between the SRM data points and the power law of the fluorescence data (Figure 6J). The live human platelet dataset was calibrated similarly using the EM data points (figure 5K). However, since EMT was performed on dehydrated, resin embedded samples, we scaled the size of EMT data points by 25% to correct for shrinkage ({Hoog:2010en}), before alignment with the live platelet fluorescence measurements.

## Discussion

Platelets are specialized cell fragments undergoing morphological changes in response to agonists, within short time scales, while in a fluid environment. We have developed a method to live image the platelet cytoskeleton while emulating similar conditions. Our technique allows precise quantification of large numbers of individual platelets while chemically perturbing cytoskeletal elements at various steps of activation. Owing to the single-cell precision of our data, we are able to address the variation in activation response using the natural heterogeneity in platelet sizes, in addition to measuring the population-scale response.

The platelet sizes we observed match those reported earlier ({Severin:2012dx}). However, our dataset offers complimentary measurements of marginal band length and intensity of microtubules in the same platelet. Both mouse and human platelets follow a similar scaling law, approximately *I*_*MT*_ ∝ *R*^2^ where R is the marginal band size (*R* = *L*_*MB*_/2π) for resting platelets) and *I*_*MT*_ is the fluorescence intensity of labeled microtubules. A different law (*I*_*MT*_ ∝ *R*^4^) was observed across species for erythrocytes with a marginal band (Dmitrieff et al., 2017), but it was already noted that platelets did not obey this scaling. The physical arguments provided to explain an exponent of 4 therefore do not hold for the data presented here, at the scale of single species. It may be that either the tension decreases with cell size, or that other factors, in particular crosslinkers and molecular motors (Diagouraga et al., 2014), enable the MB to generate more force.

Actin is known to be essential for marginal band coiling as shown previously (Diagouraga et al., 2014). Diagouraga et.al. observed that upon platelet activation with ADP, MBs coil and, on average, longer than the flat controls, which led them to propose a marginal band extension based coiling mechanism. This conclusion was however derived from comparing a set of fixed coiled cells with another set of fixed resting cells. We have seen that larger MBs are actually more prone to coiling, and suspect therefore that this approach might have suffered from a sample bias: by selecting coiled cells, they may have selected larger cells, perhaps leading to an erroneous conclusion. We saw no elongation of the marginal bands in live cells upon ADP treatment in platelets where actin was already depolymerized. On the contrary, we observe a significant shrinking of the MB length upon activation, which is too large to be an artifact of the microscopy. Our live-microscopy results clearly speaks against MB extension.

Another potential coiling mechanism could be the weakening of the marginal band by transient loss of polymerized tubulin mass upon ADP treatment as has been reported {Steiner:1979co}. However, coiling still occurs in platelets treated with taxol that prevents microtubule depolymerization {Shiba:1988tb}. In our experiments, we observe a decrease in amount of polymerized tubulin, as recorded by a relative change in marginal band intensity, upon ADP treatment of platelets (Supp. figure 3). However, the change in intensity we observe is not transient. This could indeed be attributed, either to actual depolymerization of microtubules, or to the slow labeling dynamics of our fluorescent probe, SiR tubulin, at the concentrations we use it, which could make it harder to detect newly polymerized microtubules. Additionally, we also observe no decrease in amount of polymerized tubulin, upon ADP treatment in platelets with depolymerized actin (Supp. figure 3). It suggests that microtubule depolymerization is a consequence and not the cause of marginal band coiling.

The morphological changes in platelets in response to ADP treatment are known to be reversible ({Ehrman:1978uj}, {Frojmovic:1982wk}). In our experiments, marginal bands return to a flat morphology suggesting platelets regain a discoid shape. Owing to the dynamic control of chemical perturbations in our assay, we are able to depolymerize actin during the coiling process by co-treatment with ADP and LatA. The partial response we observe in co-treatment assay results from depolymerization of actin during coiling hence affecting the maximum degree, but not the dynamics, of coiling. The recovery of MB length is complete, as we observe no lasting length reduction in the presence of LatA. This suggests that the smaller length of MBs at long times after activation, in absence of LatA, is due to persistent actin contraction.

Although it is not clearly established whether the heterogeneity in platelet size is the result of a fragmentation based manufacturing process, or an age dependent alteration of platelet size {Karpatkin:1969gc}, this variation has been demonstrated to result in different platelet response ({Corash:1977ue}{Thompson:1983vu}). Larger platelets, possibly younger, are found to be more active which is correlated to larger volume {Karpatkin:1978ut} and higher density {Corash:1977ue}. Our data shows that larger platelets achieve a higher degree of coiling, hence respond more. This can be explained taking into account the mechanics of the system, as a longer bundle of elastic rods is more likely to buckle under force as compared to a shorter one.

Electron microscopy has been used to get insight into the internal structure of platelets ({Behnke:1965wt} {White:1968tr} {ZuckerFranklin:1969ez}refs?). Some recent studies have also used electron tomography technique to study the three-dimensional internal structure of platelets {vanNispentotPannerden:2010fs} (Wang et al., 2015) {Sorrentino:2016if}. We have augmented the existing knowledge with unique information, including individual microtubule positions, numbers, lengths and polarities within the MB using electron tomography coupled with serial sectioning. We found the microtubule lengths in the MB to be exponentially distributed, which reconciles well with the previously reported presence of very long single microtubules in the MB {Nachmias:1980ws} (Kenney & Linck, 1985), approximately 100µm of polymerized tubulin, as well as presence of multiple plus and minus ends {PatelHett:2008jz}. Apart from the structural composition, an exponentially distributed microtubule population is indicative of dynamic instability, suggesting length regulation by periods of growth and shrinkage (Mitchison & Kirschner, 1984). In platelets, microtubule dynamics have been shown to mediate age dependent platelet size reduction, as treatment with paclitaxel (a drug that blocks microtubule dynamics) prevents physiological reduction in platelet size {PatelHett:2008jz}. Additionally, with our tomography measurements, we can observe a preferred center-to-center distance of around 30nm between microtubules in compact MBs. Given the microtubule diameter of around 25nm, we expect the size of potential crosslinking molecule to be around 5nm. A similar distance of approximately 5nm has been measured as an electron lucid zone surrounding microtubules {Xu:1988tb}, or even as fibrillar amorphous material of <7nm size projecting from microtubule surface (Kenney & Linck, 1985) in MBs. These distances, however, cannot be taken into account as an accurate quantitative measure as the process of high-pressure freezing followed by freeze substitution to perform electron tomography are known to induce sample shrinkage of around 25% {Hoog:2010en}. Assuming uniform shrinkage within and outside microtubules, the crosslinking distance can be up to 6.25nm. Regardless the precise value, the presence of an over-represented, preferred, distance between microtubules indicates presence of an unknown, yet specific sized cross-linking molecule.

We also used SRM to characterize the marginal band. Direct measurement of individual microtubules within the MB was not possible due to lack of resolution from the immuolabeling technique we used. This was a result of linkage error caused due to the use of a primary, anti-tubulin, antibody followed by a dye coupled secondary antibody to localize the desired epitope. This method has been shown to result in a microtubule diameter of 45.6±5.8nm {Ries:2012fa} for isolated microtubules within cells. As a result, for the MB it would not be possible to discern individual microtubules with this technique. By labeling of gamma-tubulin, we found that the localizations were distributed along the length of the MB as described before {PatelHett:2008jz}. However, the numbers of microtubules that have been reported by counting gamma tubulin foci with confocal imaging (9.06 ±1.61 foci per platelet) are significantly less than what we observed with SRM (25.2 ±6 foci per platelet). One likely reason could be that the gain of resolution in 3D gives us a clearer answer in terms of number of localizations as many confocal spots were large and possibly clusters of multiple ends. Interestingly, in activated platelets, they observed 25.5 ±8.6 EB1 (microtubule plus-end marker) comets, indicating a presence of around 25 microtubules. It suggests that most microtubules in platelet start to grow, and hence form EB1 comets, and are technically easier to mark individually as the plus ends fray out in activated platelets.

Using our experimental and analysis methods, we documented the balance of forces opposing the cortex and the marginal band. The microtubule cytoskeleton controls the size, as well as the propensity of activation of a platelet. Surprisingly, actin disruption does not lead to important MB extension, indicating a potential role for the spectrin cortex in resting platelet. It is however tempting to speculate that the compression provided by actin is necessary to assemble the spectrin cortex with a size matching the resting platelet. While investigating the role of cytoskeleton in platelet activation, we measured many important microtubules quantities, and were able to deduce how they vary with platelet size. From the data presented here, it is possible to infer the length distribution of the microtubules, their number and life times. This data will be essential to model the mechanics of platelets in details, and hopefully explain the response upon activation. As many therapies directly or indirectly target cytoskeleton mechanics, we can hope that this will lead to some important insights that would help us to better understand platelet related disorders.

